# Nutrient sensing in the nucleus of the solitary tract mediates non-aversive suppression of feeding via inhibition of AgRP neurons

**DOI:** 10.1101/2020.06.23.167494

**Authors:** Anthony H. Tsang, Danae Nuzacci, Tamana Darwish, Havish Samudrala, Clémence Blouet

## Abstract

The nucleus of the solitary tract (NTS) is emerging as a major site of action for the appetite-suppressive effects of leading pharmacotherapies currently investigated for the treatment of obesity. However, our understanding of how NTS neurons regulate appetite remains incomplete. Here we used NTS nutrient sensing as an entry point to characterize stimulus-defined neuronal ensembles engaged by the NTS to produce physiological satiety. Using activity-dependent expression of genetically-encoded circuit analysis tools, we found that NTS detection of leucine engages NTS prolactin-releasing peptide (PrRP) neurons to inhibit AgRP neurons via a population of leptin-receptor-expressing neurons in the dorsomedial hypothalamus. This circuit is necessary for the anorectic response to NTS leucine, the appetite-suppressive effect of high protein diets, and the long-term control of energy balance. These results extends the integrative capability of AgRP neurons to include brainstem nutrient sensing inputs.

## Introduction

The nucleus of the solitary tract (NTS) is established as a major brain site for the sensing and integration of signals relevant to the control of feeding behavior. It is a neuroanatomical hub for ascending vagal afferents activated by ingested foods, corticolimbic-descending inputs encoding homeostatic, cognitive and motivational controls of feeding, and blood-borne signals diffusing from the adjacent area postrema (AP) that lacks a blood-brain barrier (Grill & Hayes, 2009). Molecularly, it is enriched in specialized interoceptive neuronal populations equipped to monitor circulating levels of nutrients, gut hormones and adiposity signals (Blouet & Schwartz, 2012).

NTS processing of these diverse inputs is classically described as the main mediator of the short-term negative feedback control of ingestion (or satiation) via recruitment of medullary motor output circuits (Grill & Hayes, 2009). The NTS also relays processed information to the lateral parabrachial nucleus (lPBN), established as a common target for NTS efferents in the central representation of aversive and avoidance feeding-related cues (D.-Y. Kim et al., 2020; Palmiter, 2018). In both cases, the NTS outputs interrupts food ingestion, and until recently had not been implicated in the regulation of hunger, the long-term control of satiety, or hedonic feeding. Studies applying molecular genetics or modern circuit analysis tools to the functional characterization of NTS neurons revealed that the NTS can in fact modulate a much bigger range of behavioral effectors of energy balance including meal initiation and satiety (Blouet & Schwartz, 2012; D’Agostino et al., 2016; Gaykema et al., 2017; Hayes et al., 2011; Su et al., 2017). Yet, little is known about the neural mechanisms through which the NTS regulates forebrain hunger and satiety circuits, and the physiological contexts in which these NTS feeding-regulatory forebrain-projecting outputs are engaged.

Conceptually, a key question is whether different behavioral effectors of ingestion (i.e. satiation, avoidance/aversion, satiety, food-seeking) are engaged by distinct and functionally specialized NTS neuronal subsets. Evidence that segregated subsets of CCK^NTS^ or TH^NTS^ neurons project to the lPBN or hypothalamus to produce either avoidance/aversive anorexia vs. satiety or glucoprivic feeding support this view (Aklan et al., 2020; Roman et al., 2017). Alternatively, or in addition to this possibility, recruitment of the same neurons could simultaneously or gradually evoke multiple of these behavioral outputs. The fact that NTS catecholaminergic neurons send collaterals to midbrain and forebrain targets provides a neuroanatomical basis for the latter (Petrov et al., 1993), which could explain the ability of high doses of the satiation hormones like CCK to recruit aversive circuits (Swerdlow et al., 1983), and/or provide a mechanism for the synergistic feeding suppressive effects produced by the combination of anorectic signals (Bhavsar et al., 1998; Blevins et al., 2009; Blouet & Schwartz, 2012). Addressing this question with molecularly-defined circuit analysis tools is difficult because most identified NTS neurochemical subsets are functionally heterogeneous, respond to multiple cues and project widely throughout the neuraxis (D’Agostino et al., 2016; Rinaman, 2010). Instead, it may be possible to better understand the functional organization of NTS feeding-regulatory circuits using functionally-defined circuit mapping, which could be particularly insightful if subsets of NTS neurons are specialized in the transmission of highly specific sensory cues and organized in a similar fashion as gustatory and vagal sensory neurons (Bai et al., 2019; Williams et al., 2016). Applying such a strategy to signals able to produce satiation or satiety without negative consequences may lead to important new understanding of how to pharmacologically suppress appetite without undesirable side effects.

We previously showed that NTS sensing of the branched-chain amino acid leucine not only modulates the control of meal size but also rapidly suppresses hunger in fasted animals and increases satiety without the production of conditioned taste aversion (Blouet & Schwartz, 2010; Cheng et al., 2020). Here, we used NTS leucine sensing as a functional entry point to investigate ascending neural circuits engaged by NTS neurons to modulate hunger and satiety. In these experiments, Leucine is injected into the NTS at physiologically relevant doses to model the postprandial increase in brain leucine levels as seen in response to the consumption of high-protein meals, a dietary paradigm that potently suppresses food intake. We employed an activity-dependent labelling and circuit mapping strategy which allowed to express circuit analysis tools specifically in leucine-sensing neurons and downstream circuits.

## RESULTS

### NTS amino acid sensing inhibits AgRP neurons via a polysynaptic circuit

NTS leucine sensing rapidly reduces hunger in fasted rodents(Blouet & Schwartz, 2012; Cavanaugh et al., 2015), but the underlying neural circuit mediating this response in unknown. To characterize the ascending neural circuits engaged by NTS leucine sensing to rapidly inhibit appetite, we first assessed neuronal activation throughout the neuraxis in response to local bilateral NTS leucine administration. Mice were fasted for 6h and received a site-specific injection of 50nl of leucine per side into the caudomedial NTS as previously described (Cavanaugh et al., 2015) (Fig. 1a). Neuronal activity was assessed using c-fos immunohistochemistry 80 min later. NTS leucine induced robust c-fos expression in the caudomedial NTS as well as in the adjacent area postrema (AP) (Fig. 1b, 1c). Outside this region, only a few brain sites were significantly activated by local NTS leucine administration compared to aCSF vehicle: the locus coeruleus (LC), and the paraventricular, ventromedial and dorsomedial nuclei of the hypothalamus (Fig. 1b, 1c). In contrast, NTS leucine produced a 2-fold decrease in c-fos immunolabelling in the ARH (Fig. 1b, 1c). Of note, NTS leucine did not produce neuronal activation in the parabrachial nucleus (PBN, Fig. 1b), consistent with the lack of conditioned avoidance in response to parenchymal NTS leucine administration in mice (Cheng et al., 2020).

**Fig 1:**
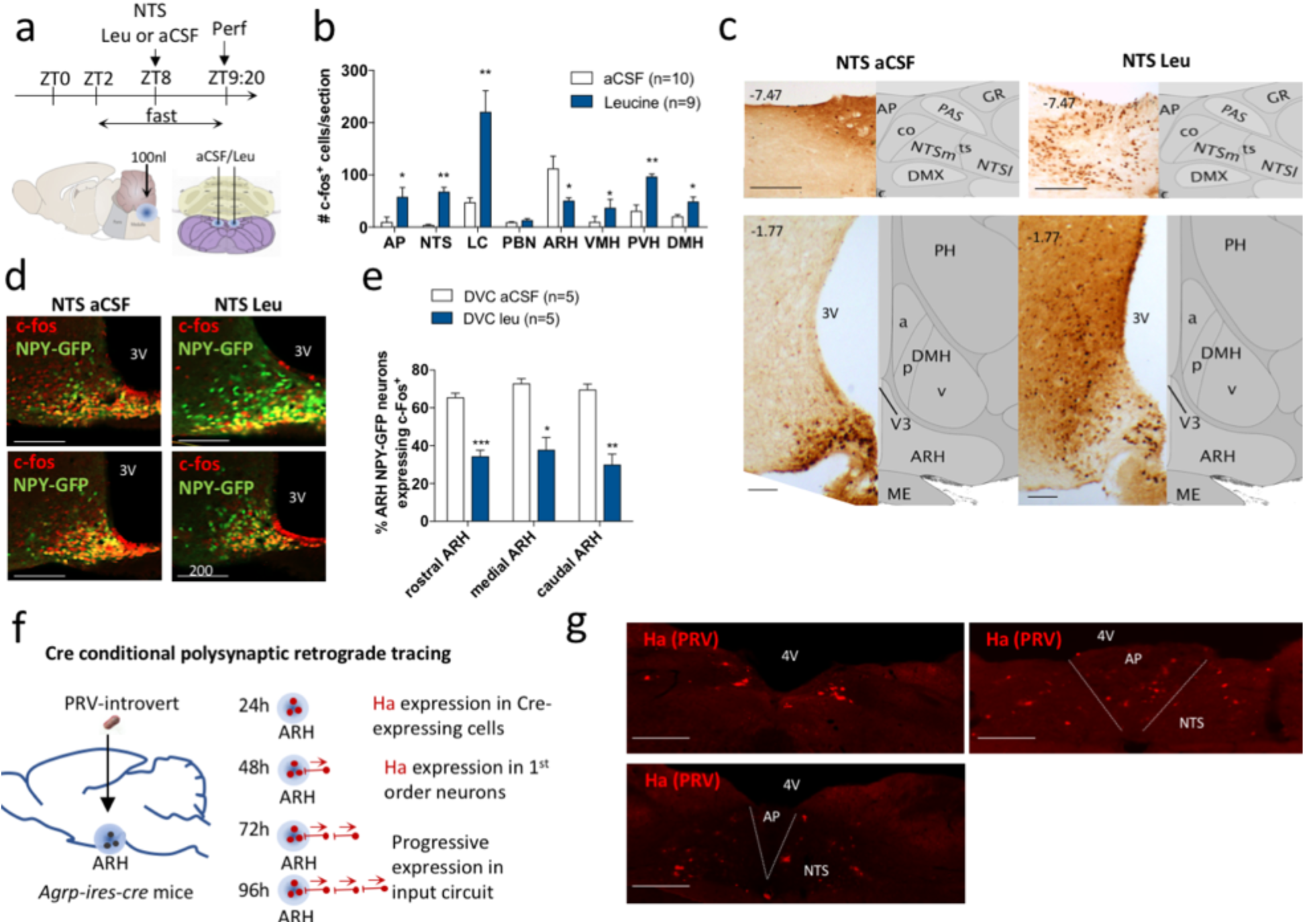
NTS amino acid sensing inhibits AGRP neurons via a polysynaptic circuit. Injection paradigm (a) used to activate NTS leucine-sensing neural circuits in mice. Quantification of c-fos immunolabelling (b) and representative images (c) from selected sites of the mouse brain following NTS leucine delivery. Representative images (d) and quantification (e) of c-fos immunodetection in AgRP/NPY neurons in the ARH. Protocol for Cre conditional polysynaptic retrograde tracing using PRV-Introvert (f) and expression of the PRV-Introvert reporter Ha in the caudomedial NTS after 96h (g). Scale bar =200um. AP: area postrema, NTS: nucleus of the solitary tract, LC: locus coereleus, PBN: parabrachial nucleus, ARH: arcuate nucleus of the hypothalamus, VMH: ventromedial nucleus of the hypothalamus, PVH: paraventricular nucleus of the hypothalamus, DMH: dorsomedial nucleus of the hypothalamus, 3V : 3^rd^ ventricle, 4V: 4^th^ ventricle. Rostral ARH: Bregma -1.07 to -1.37, Medial ARH: Bregma -1.37 to -1.77, Caudal ARH: Bregma -1/77 to-2.07. *: p<0.05 vs. aCSF; **: p<0.01 vs. aCSF; ***: p<0.001 vs. aCSF. All results are shown as means ± SEM.

**Supp. Fig. 1:**
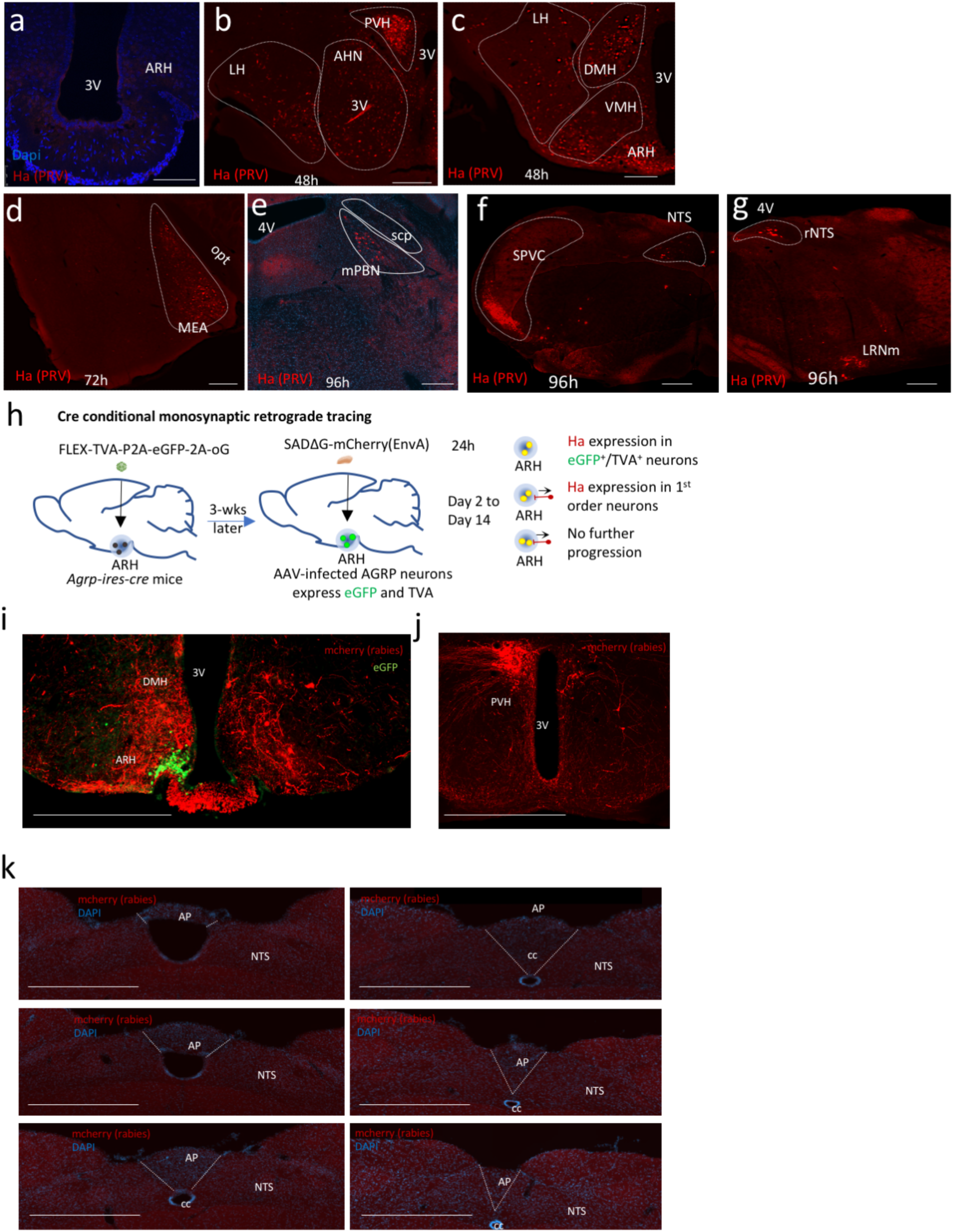
Cre-dependent retrograde polysynaptic and monosynaptic viral tracing in *Agrp-ires-cre* mice. HA immunodetection in WT mice 96h post inoculation (a) and in *Agrp-ires-cre* mice 48h (b, c) 72h (d) and 96h (e, f, g) after a bilateral injection of PRV-Introvert in the ARH (Scale bar: 400um). Protocol for Cre conditional monosynaptic retrograde tracing using SADΔG-mCherry(EnvA) (h) and expression of the mCherry and eGFP in the hypothalamus (i, j) and hindbrain (k) of *Agrp-ires-cre* mice unilaterally infected with rAAV8-hSyn-FLEX-TVA-P2A-eGFP-2A-oG and SADΔG-mCherry(EnvA) 2-weeks after rabies infection (Scale bar: 800um). 3V: 3rd ventricle, ARH: arcuate nucleus of the hypothalamus, PVH: paraventricular hypothalamic nucleus, DMH: dorsomedial hypothalamic nucleus, LH: lateral nucleus of the hypothalamus, VMH: ventromedial hypothalamic nucleus, opt: optical tract, MEA: medial amygdala, mPBN: medial parabrachial nucleus, scp: superior cerebelar peduncles, AHN: anterior hypothalamic nucleus, NTS: nucleus of the solitary tract, AP: area postrema, cc: central canal, rNTS: rostral nucleus of the solitary tract, SPVC: Spinal nucleus of the trigeminal, LRNm: Lateral reticular nucleus, magnocellular part.

The ARH contains intermingled orexigenic and anorexigenic neurons including AgRP neurons, critical for the development of food seeking behavior and meal initiation in hungry mice (Atasoy et al., 2012; Fenselau et al., 2017). We previously found that NTS leucine sensing robustly increases first-meal latency in fasted mice, hence reduces the drive to approach and consume food (Blouet & Schwartz, 2012); (Cavanaugh et al., 2015). This, together with the reduced c-fos expression in the ARH following NTS leucine administration, prompted us to hypothesize that hindbrain leucine sensing may rapidly inhibit AgRP neurons. To test this, we repeated the same experiment (Fig. 1a) in *Npy-hrGFP* transgenic mice, where the hrGFP signal in the mediobasal hypothalamus selectively labels all AgRP neurons (Hahn et al., 1998). A majority of ARH NPY/AgRP neurons were activated under control conditions (Fig. 1d, 1e). As predicted, NTS leucine produced a 2-fold decrease in NPY/AgRP neuronal activation throughout the rostro-caudal extend of the ARH (Fig. 1d, 1e). Thus, NTS leucine sensing rapidly inhibits ARH NPY/AgRP neurons.

Previous work indicates that NTS inputs can modulate the activity of AgRP (Aklan et al., 2020; Cheng et al., 2020), but the neuroanatomical organization of these inputs and the physiological conditions in which they are engaged to modulate feeding remain unclear. To establish that AgRP neurons are synaptically connected to NTS neurons we performed a series of retrograde viral tracing studies. Pseudorabies virus Bartha strain (PRV-Bartha) is a neuroanatomical tracer that is transmitted retrogradely across synapses and can be used to define polysynaptic inputs to infected neurons (Pickard et al., 2002). PRV-Introvert is a newly developed version of PRV-Bartha in which retrograde viral propagation and reporter expression are activated only after exposure to Cre recombinase with high specificity (Pomeranz et al., 2017). We used PRV Introvert in *Agrp-ires-cre* mice to serially label chains of presynaptic neurons projecting to ARH AgRP neurons. Mice were sacrificed 0, 24, 48, 72h or 96h after local ARH inoculation with PRV-Introvert, and brains were examined for HA reporter expression (Fig. 1f). As expected, we did not detect any HA immunolabelling in the brain of wild type mice injected with the virus (Suppl 1a)., confirming cre-dependency. At 24 h after injection, PRV-Introvert was detectable in the ARH, indicating that cre-mediated recombination occurred locally within 24 h of PRV injection. After 48h, spread of the cre-activated PRV virus was observed in multiple hypothalamic sites including the arcuate, ventromedial, dorsomedial, lateral and paraventricular nuclei (Suppl. 1b, 1c). After 72h, the medial amygdala was labelled (Suppl. 1d). After 96h, we detected PRV in several pontine, midbrain and hindbrain structures. These included the medial parabrachial nucleus (mPBN) (Suppl Fig. 1e), the ventrocaudal part of the spinal trigeminal nucleus (vcSPVC) (Suppl Fig. 1f), the rostroventrolateral medulla (RVLM) (Supp. Fig. 1e), the AP and the NTS, both in its rostral portion and in the lateral portion of the caudomedial NTS (Fig 1g, Suppl Fig 1f-1g). Thus, ARH AgRP neurons receive inputs from multiple midbrain and hindbrain sites, including the caudomedial NTS.

The long survival time necessary to detect the presence of PRV in the NTS suggests that the NTS→ AgRP circuit contains more than 1 synapse. Alternatively, the long distance that the virus needs to travel to label hindbrain sites may also explain the lack of signal at 48 and 72h. To clarify this, we performed cre-dependent monosynaptic retrograde viral tracing in *Agrp-ires-cre* mice using an envelope protein (EnvA) pseudotyped glycoprotein (g)-deleted rabies virus modified to express mCherry (SADΔG-mCherry(EnvA) (Callaway & Luo, 2015; E. J. Kim et al., 2016). AgRP-cre expressing neurons were first genetically modified to co-express TVA (receptor for the avian sarcoma leucosis virus glycoprotein EnvA) and oG (optimized rabies envelope glycoprotein) via targeted unilateral injections of rAAV8-hSyn-FLEX-TVA-P2A-eGFP-2A-oG into the ARH (Suppl. Fig. 1h). AgRP neurons infected with this construct became selectively competent for transduction by SADΔG-mCherry(EnvA) and expressed eGFP. 3 weeks later, mice received a unilateral injection of SADΔG-mCherry(EnvA) in the same injection site, and were sacrificed at various survival times (up to 14 days). Brains were processed to examine mcherry expression. Two weeks after the SADΔG-mCherry(EnvA) injection, we observed dense eGFP expression in the ARH (Suppl. Fig. 1i) together with dense mCherry immunolabelling in the ARH, DMH and PVH (Suppl. Fig. 1i, 1j). 42% of AgRP neurons expressing eGFP co-expressed mCherry. We carefully examined the NTS of 8 successfully infected animals throughout the rostro-caudal extend of the NTS but did not detect rabies-infected cell bodies (Suppl. Fig. 1k). These data support the conclusion that the NTS does not send monosynaptic inputs to AgRP neurons.

### NTS PrRP neurons are leucine-sensing and project to AgRP^ARH^ neurons

Previous work showed that a majority of NTS leucine-sensing neurons express TH (Blouet & Schwartz, 2012; Cavanaugh et al., 2015), but TH labels a molecularly and functionally diverse group of neurons, prompting us to further analyze the neurochemical identity of NTS neurons responsive to leucine. The NTS contains several neuronal subpopulations responsive to aversive gastrointestinal stimuli or nutritional stress, leading to the formation of visceral malaise, taste aversion or avoidance (Callaway & Luo, 2015; Holt et al., 2019; E. J. Kim et al., 2016; Patel et al., 2019; Roman et al., 2016). In contrast, some NTS neuronal subtypes are recruited preferentially in response to physiological satiation cues and do not produce aversive anorexia even in the context of pharmacological activation. These include subsets of TH^NTS^ neurons expressing prolactin-releasing peptide (PrRP) (Kreisler et al., 2014; Lawrence et al., 2002), and recently characterized Calcr^NTS^ neurons (Cheng et al., 2020). We previously showed that NTS leucine does not produce conditioned avoidance (Cheng et al., 2020), leading us to hypothesize that leucine specifically engages either PrRP^NTS^ or Calcr^NTS^ neurons to suppress feeding. To examine this possibility, we used RNAscope multiplex in situ hybridization (ISH) against *Fos, Prlh* (transcript for PrRP) *and Calcr* in caudomedial hindbrain sections of mice that received local NTS leucine injections as described above (Fig. 1a). We found a significant overlap between PrRP^NTS^ and Calcr^NTS^ neurons (Fig. 2a): 75.1±1.3% PrRP^NTS^ neurons expressed *Calcr*, while 68.3±0.9% Calcr^NTS^ neurons expressed *Prlh*. Most of NTS *Calcr*^+^*/ Prlh*^-^ neurons where concentrated in the dorsal NTS, and *Calcr* was also expressed in a dense *Prlh*^-^ neuronal population in the AP (Fig 2a). Consistent with c-fos immunolabelling, we found that *Fos* expression rapidly increased in the NTS and AP in response to local leucine delivery (Fig. 2a-2b). Leucine activated on average 80% of PrRP^NTS^ neurons (Fig 2a, 2c) which represented 34.7±8.6% of the total population activated by leucine in the caudomedial NTS. Analysis of high content ISH images revealed that the number of PrRP^NTS^ neurons was similar between conditions (Fig. 2d), but leucine increased the expression of *Prlh* (Fig. 2e), introducing a role for PrRP neurotransmission in leucine-sensing neurocircuits.

**Figure 2:**
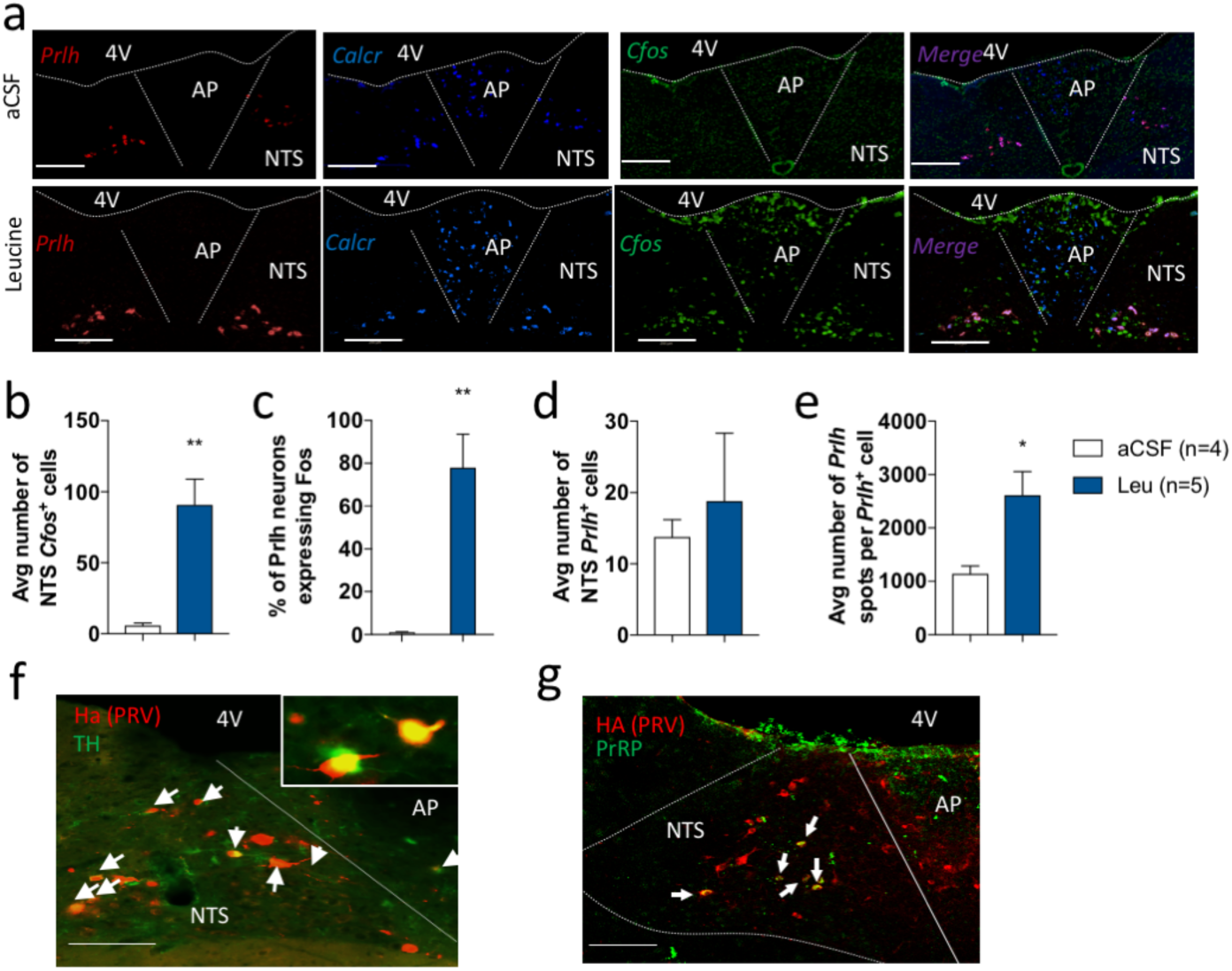
Neurochemical characterization of NTS leucine-sensing neurons. Representative images (a). and quantification (b-e) of multiplexed in situ hybridization against *Prlh, Calcr* and *Cfos* in the dorsovagal complex of mice after an injection of leucine into the NTS. Representative images of the colocalization of the Ha reporter from the polysynaptic retrograde virus PRV-Introvert and tyrosine hydroxylase (TH, f) or prolactin releasing peptide (PrRP, g) by multiplexed immunofluorescent labelling in the NTS of *Agrp-ires-cre* mice 96h after PRV-Introvert delivery into the ARH. Scale bar: 200um. 4V: 4^th^ ventricle. AP: area postrema, NTS: nucleus of the solitary tract. *: p<0.05 vs. aCSF; **: p<0.01 vs. aCSF; ***: p<0.001 vs. aCSF. All results are shown as means ± SEM.

**Supp. Fig. 2:**
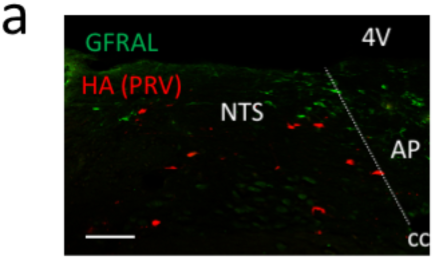
GFRAL and HA immunolabelling in the NTS of *Agrp-ires-cre* mice 96h after PRV-Introvert delivery into the ARH. Scale bar: 200um. 4V: 4^th^ ventricle. AP: area postrema, NTS: nucleus of the solitary tract.

We then determined whether PrRP^NTS^ neurons project to ARH^AgRP^ neurons using brain sections from *Agrp-ires-cre* mice infected with PRV-Introvert and killed 96h after infection. We found that a majority of the HA^+^ neurons labelled 96 h after ARH PRV-Introvert delivery colocalized with TH (61±8%) and PrRP (35±2%) (Fig. 2f, 2g) confirming that PrRP^NTS^ neurons project to ARH^AgRP^ neurons. Of note, NTS HA^+^ neurons did not express GDF15 receptor GFRAL (Suppl. Fig 2a), indicating that GDF15 does not engage the NTS→ AgRP^ARH^ circuit to suppress feeding.

### NTS leucine sensing activates DMH LepR^+^/GPR10 neurons projecting to AgRP neurons

We next investigated the neuronal populations relaying NTS leucine-sensing inputs to AgRP neurons. Given that PrRP^NTS^ neurons represent only a third of NTS leucine-sensing neurons and a third of NTS neurons projection to AgRP^ARH^, and in the absence of a known specific molecular marker for NTS leucine-sensing neurons, we developed a strategy of activity-dependent circuit mapping following NTS leucine administration. We use the AAV8-Fos-ERT2-Cre-ERT2-PEST (AAV-Fos-CreERT2) virus to translate temporally delimited neuronal activity into sustained reporter expression (Ye et al., 2016). Neurons expressing AAV-Fos-CreERT2 do not express Cre unless acutely exposed to an activating stimulus together with tamoxifen. Low dose of the tamoxifen metabolite 4-hydroxytamoxifen (4-OHT) allows genetic labelling of transiently activated neurons with high temporal specificity and low background (Ye et al., 2016). *Npy-hrGFP* mice received a co-injection of AAV-Fos-CreERT2 and AAV8-EF1a-DIO-hChR2(H134R)-mCherry viruses into the caudomedial NTS. 3 weeks later, mice were exposed to 3 experimental inductions, each separated by 96h (Fig. 3a). During each of these, mice were fasted for 4h during the light phase and received a bilateral NTS injection of aCSF or leucine followed 80 min later by an ip injection of 40mg/kg 4-OHT (hereafter designated as aCSF_induced_ and Leu_induced_ mice, respectively). Access to food was restored 4h later to avoid food-induced Cre recombination. Mice were sacrificed 14 days after the last NTS injection. Using mCherry immunodetection in brain tissues, we characterized the neuroanatomical distribution of axonal projections and synaptic terminals of NTS leucine sensing neurons. mCherry expression was dense in the caudomedial NTS of mice induced with NTS leucine injections, confirming the success of the approach (Fig. 3b). We did not detect mCherry^+^ signal in the ARH of Leu_induced_ mice, indicating that NTS leucine-sensing neurons do not project directly to the ARH (Fig. 3c). In contrast, we found mCherry^+^ fibers and terminals in the PVH and the ventral DMH of Leu_induced_ mice compared to controls (Fig. 3d-e). Thus, NTS leucine-sensing neurons project to the PVH and the DMH.

**Figure 3:**
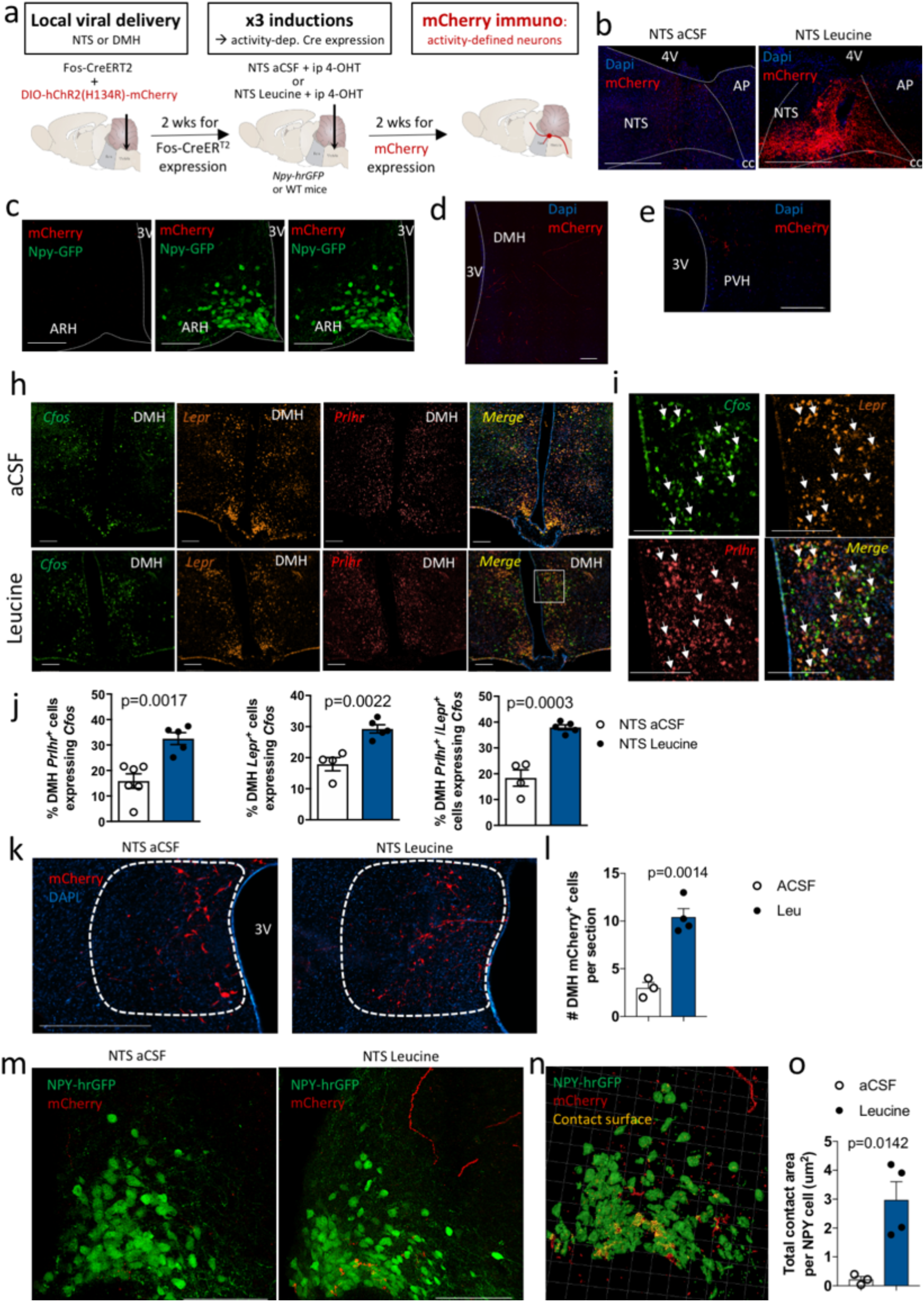
NTS leucine sensing activates DMH LepR^+^/GPR10 neurons projecting to AgRP neurons. Diagram of experimental paradigm (a) and representative images of mCherry immunolabelling in the DVC (b), ARH (c), DMH (d) and PVH (e) for activity-dependent mapping of projection outputs of NTS leucine-sensing neurons using AAV-Fos-CreERT2 and AAV8-EF1a-DIO-hChR2(H134R)-mCherry viruses. Representative images (h), high magnification images (i) and quantifications (j) of the expression of *Fos in Lepr*^+^ and *Prlhr*^+^ neurons in the DMH of mice following an injection of aCSF or leucine in the NTS. Representative images (k,m), quantification (l,o) and IMARIS 3-d reconstruction (n) of mcherry immunolabelling in the DMH (k-l) and ARH (m-o) of *Npy-hr-GFP* mice following injection of AAV-Fos-CreERT2 and AAV8-EF1a-DIO-hChR2(H134R)-mCherry viruses in the DMH and inductions with NTS aCSF or leucine. Scale bar is 200um. All results are shown as means ± SEM.

**Supp. Fig. 3:**
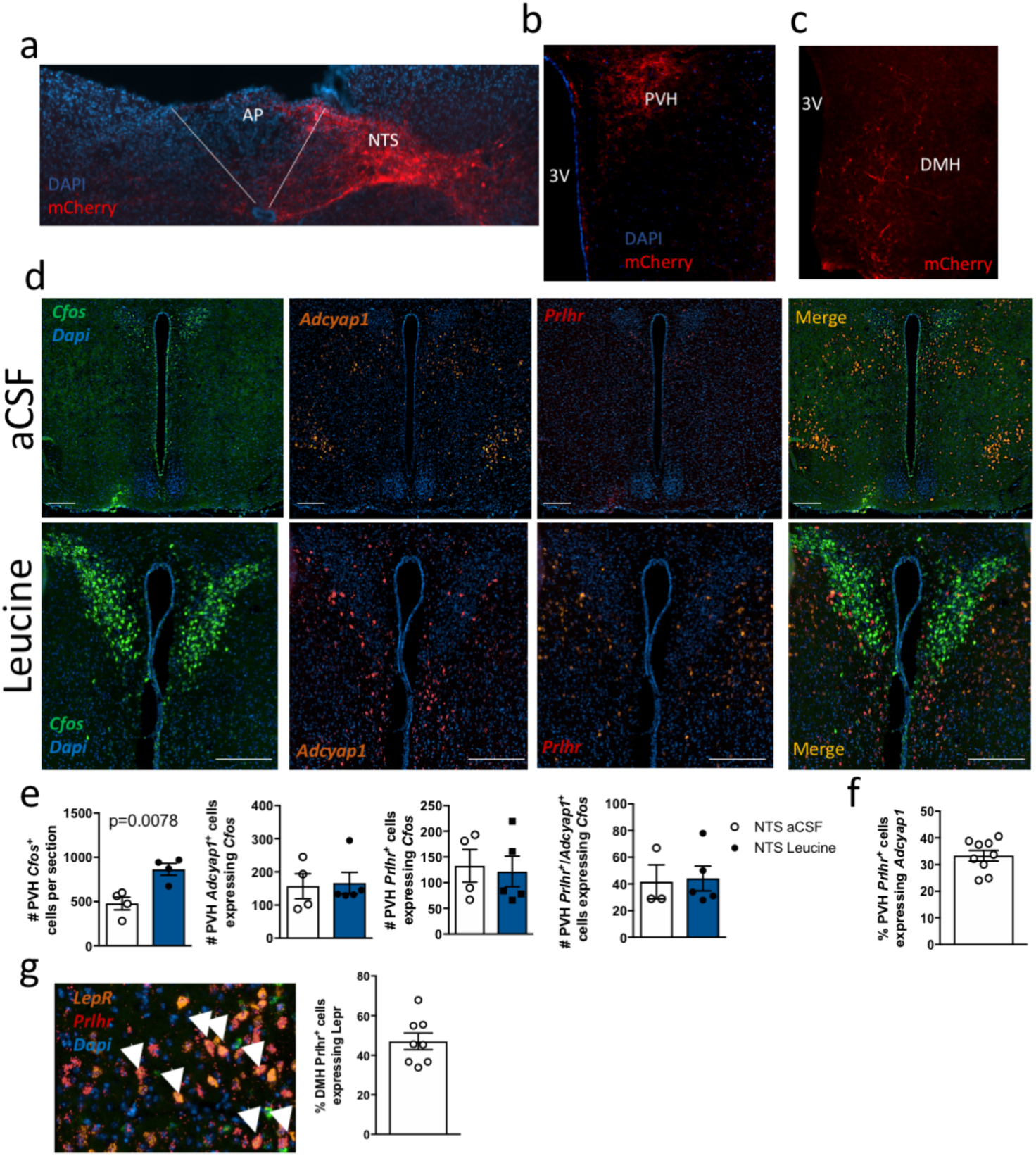
Representative images on mcherry expression in the NTS (a), PVH (b) and DMH (c) in *Th-cre* mice that received a unilateral injection of AAV8-EF1a-DIO-hChR2(H134R)-mCherry into the NTS to label synaptic terminals of NTS TH neurons. Representative images (d) and quantification (e) of *Cfos, Adcyap1* and *Prlhr* expression in the PVH of mice exposed to NTS aCSF or Leucine. Quantification of *Prlhr* and *Adcyap1* co-expression in the PVH (f). Representative image and quantification (g) of Prlhr and *LepR* co-expression in the DMH. Scale bar is 200um. All results are shown as means ± SEM.

The PVH and DMH are both good candidates to relay NTS leucine-sensing inputs from PrRP^NTS^ neurons to AgRP neurons. Both the PVH and the DMH receive dense projections from TH^NTS^ neurons (Suppl. Fig 3a-3c), are innervated by PrRP^+^ fibers, and express GPR10, the receptor for PrRP (Dodd & Luckman, 2014). Previous monosynaptic retrograde tracing studies identified the PVH and the DMH as the main sources of pre-synaptic inputs to AgRP neurons (Krashes et al., 2014), and channelrhodopsin-assisted circuit mapping studies showed that all PACAP^PVH^ and LepR^DMH^ neurons project to and directly regulate the activity of AgRP neurons (Garfield et al., 2016; Krashes et al., 2014). However, there is limited understanding of how these neuronal inputs to AgRP neurons may be engaged under physiological conditions to modulate appetite. To examine the role of PACAP^PVH^ and LepR^DMH^ in relaying leucine-sensing information from the NTS to AgRP neurons, we first used RNAScope to colocalize *Fos, Adcyap1* (transcript for PACAP) and *Prlhr* (transcript for PrRP receptor) or *Fos, LepR* and *Gpr10* in the PVH and DMH respectively of mice that received NTS aCSF or leucine as previously described (Fig. 1a). These experiments confirmed that NTS leucine produces a significant increase in the number of *Fos-*expressing neurons in the PVH compared to vehicle injection (Suppl. Fig. 3d) but NTS leucine did not produce an increase in the number of *Prlhr*^+^, *Adcyap1* ^+^ or *Prlhr*^+^/*Adcyap1* ^+^ PVH neurons expressing *Fos* (Suppl. Fig. 3d-3e). In the DMH, NTS leucine increased *Fos* expression in a group of neurons concentrated in the caudal DMH (fig. 3h). 30% of DMH neurons co-expressed *LepR* and *Prlhr* (Suppl. Fig. 3g), and NTS leucine significantly increased the number of DMH *Prlhr*^+^, *Lepr*^+^ and *Lepr*^+^/*Prlhr*^+^ neurons expressing *Fos (*Fig. 3i-3j). Thus, NTS leucine activates neurons in the DMH that are well positioned to receive inputs from NTS PrRP neurons and project to AgRP neurons.

To confirm that the DMH relays NTS leucine sensing inputs to AgRP neurons, we performed activity-dependent circuit mapping from DMH neurons activated by NTS leucine. We delivered AAV-Fos-CreERT2 and AAV8-EF1a-DIO-hChR2(H134R)-mCherry viruses into the DMH of *NPY-hrGFP* mice and exposed mice to the same induction paradigm as above (fig. 3a) to label axons and synaptic terminals of DMH neurons activated by NTS leucine. In the presence of 4-OHT, NTS leucine induced a significant increase in the number of neuronal cell bodies labelled with mCherry in the DMH (Fig 4k, 4l), confirming the success of the approach to label DMH neurons responsive to NTS leucine. We observed mcherry-labelled axons in the ARH (Fig. 3m) but failed to identify additional mcherry labelling in other brain regions (not shown). These data indicate that DMH neurons activated by NTS leucine provide axo-somatic innervation of the ARH. The projection field labelled with mCherry overlapped with the hrGFP immunofluorescent labelling of NPY/AgRP neurons (Fig. 3m). Analysis of mCherry^+^ puncta contacting ARH NPY-GFP neurons confirmed that DMH leucine-sensing neurons innervate AgRP neurons (Fig. 3n-3o). Thus, NTS leucine activates DMH neurons that project to AgRP neurons.

**Figure 4.**
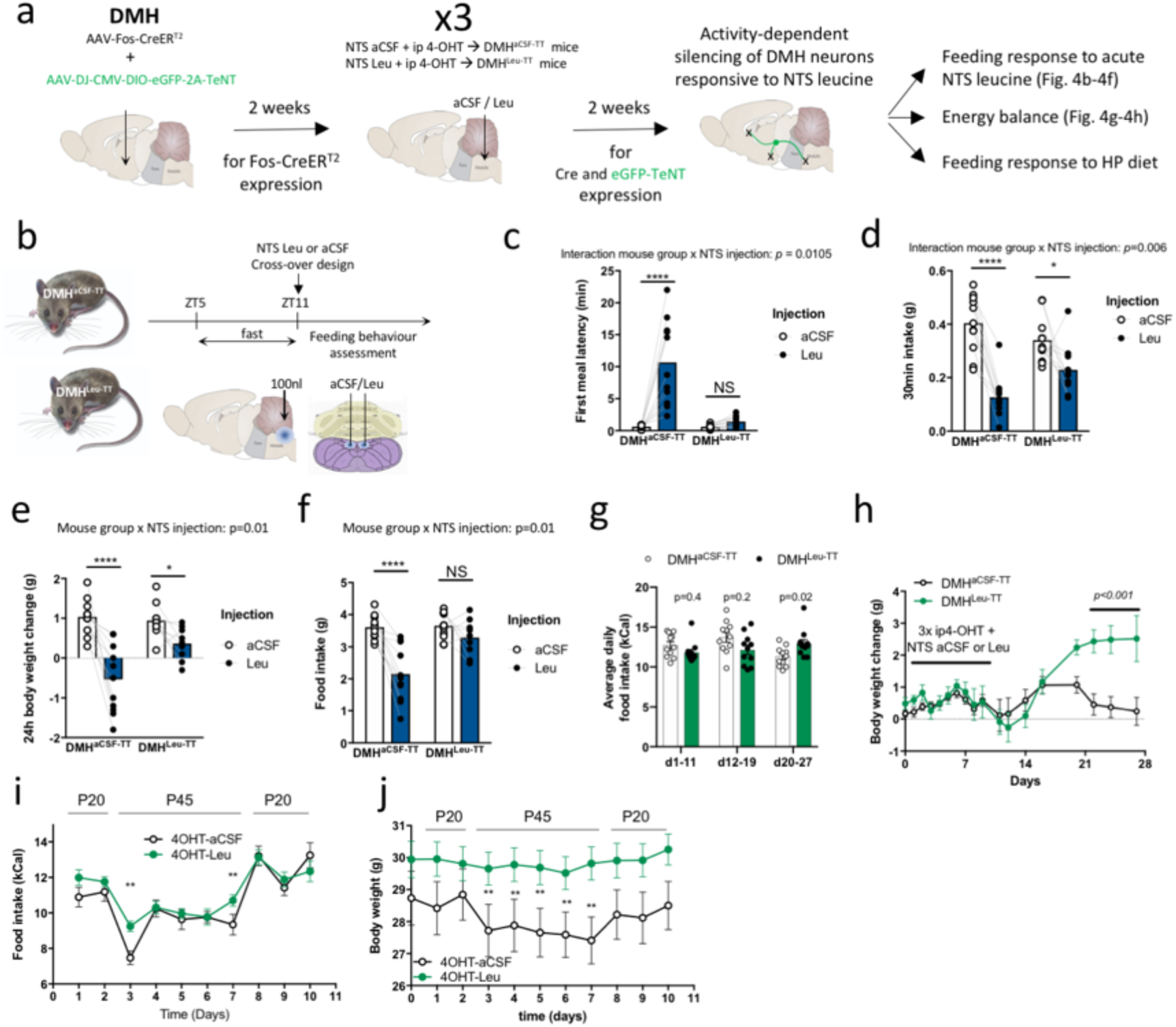
DMH neurons responsive to NTS leucine sensing are necessary for the anorectic effect of NTS leucine and high protein diets. Diagram of the experimental paradigm to selectively silence DMH neurons receiving inputs from NTS leucine-sensing neurons (a, DMH^aCSF-TT^ and DMH^leu-TT^ mice). Diagram of the experimental paradigm used to test the feeding effect of NTS leucine in DMH^aCSF-TT^ and DMH^leu-TT^ mice (b). First meal latency (c), first meal size (d) 24h body weight change (e) and 24h food intake (f) in DMH^aCSF-TT^ and DMH^leu-TT^ mice following an acute injection of aCSF or leucine into the NTS. Average food intake (g) and weight change (h) of in DMH^aCSF-TT^ and DMH^leu-TT^ mice during the 4 weeks following DMH injection with the Tet-Tox virus. Average food intake (i) and body weight (j) in DMH^aCSF-TT^ and DMH^leu-TT^ mice during transitions from diets containing 20% or 45 % of energy as proteins. All results are shown as means ± SEM. *: p<0.05, **: p<0.01, ***: p<0.001 and ***:p<0.0001 vs. aCSF or control group.

**Supp. Fig. 4:**
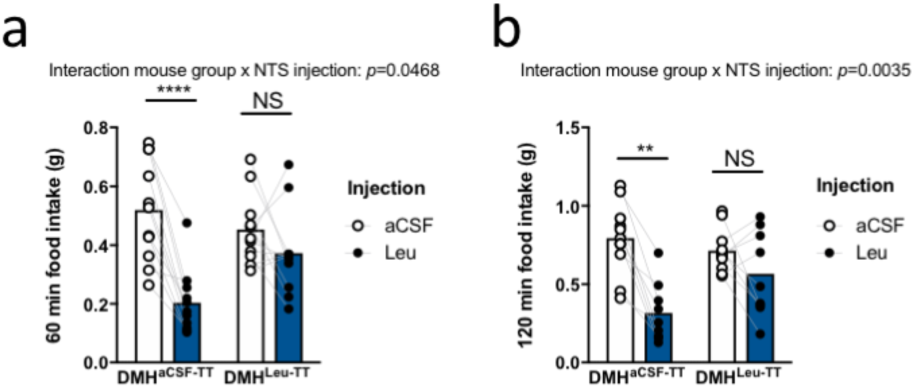
60min (a) and 120 min (b) food intake in DMH^aCSF-TT^ and DMH^leu-TT^ mice following an acute injection of aCSF or leucine into the NTS.

### DMH neurons responsive to NTS leucine sensing are necessary for the anorectic effect of NTS leucine and high protein diets and the long-term control of energy balance

To directly test the role of DMH neurons engaged downstream from NTS leucine sensing in the appetite-suppressing effect of NTS leucine, we selectively silenced DMH neurons activated by NTS leucine using cell-specific expression of tetanus toxin (TT) to prevent synaptic neurotransmitter release. To achieve this, we co-injected AAV-Fos-CreERT2 and AAV-DJ-CMV-DIO-eGFP-2A-TeNT into the DMH of wild-type mice and exposed to the same induction paradigm as above via NTS injections of aCSF or leucine (DMH^aCSF-TT^ and DMH^leu-TT,^ respectively) (fig. 4a). We then compared the anorectic response to NTS leucine in DMH^leu-TT^ and DMH^aCSF-TT^ mice (Fig. 4b). In DMH^aCSF-TT^ controls, NTS leucine produced the expected behavioral response, including an increase meal latency and decrease in food intake following the injection (Fig. 4c, 4d, Suppl movie 1, Suppl. Figure 4a, 4b). In contrast, NTS leucine failed to increase meal latency and decrease food intake in DMH^leu-TT^ mice (Fig. 4c, 4d, Suppl movie 2, Suppl. Figure 4a, 4b). Thus, DMH neurons engaged by NTS leucine sensing are required for the acute effects of NTS leucine on meal initiation and satiation. In addition, NTS leucine-induced reductions in 24h food intake and 24h weight change were blunted in DMH^leu-TT^ mice (Fig. 4e, 4f), supporting a role for the NTS→DMH leucine-sensing circuit in the long-term feeding and metabolic consequences of NTS leucine sensing. Of note, over time, DMH^Leu-TT^ developed a slight but significant hyperphagia (Fig. 4g) and gained significantly more weight than the DMH^aCSF-TT^ controls (Fig. 4h), supporting a role for DMH neurons receiving NTS leucine-sensing inputs in the chronic maintenance of energy balance.

Last, we asked whether this newly characterized circuit was relevant not only for the feeding-suppressive effect of NTS leucine, but also for the anorectic response to high-protein feeding. In mice, acute exposure to a high protein diet reduces appetite and weight gain (Vu et al., 2017). While the central mechanisms mediating these responses are poorly characterized, the NTS is established a neuroanatomical site responding to high protein diets (Darcel et al., 2005). Furthermore, a single high-protein meal is sufficient to increase brain leucine concentration (Darcel et al., 2005), supporting the possibility that NTS leucine -sensing neurons could mediate appetite suppression in response to dietary proteins. To address this, we exposed DMH^Leu-TT^ and DMH^aCSF-TT^ mice to a high protein diet containing 45% of energy as protein (P45) and isocaloric with the control maintenance diet containing 20% energy as protein (P20, control maintenance diet). To avoid a neophobic response to the P45 diet, mice were first briefly exposed to P45 pellets (3 times for 30-min on 3 consecutive days). A week later, mice were switched to the P45 diet for 6 days. In control DMH^aCSF-TT^ mice, the P45 diet produced a rapid 30% decrease in energy intake, followed by a sustained 10 to 15% reduction in daily energy intake in the following days (Fig. 4i). The anorectic response to the high protein diet was associated with sustained weight loss (Fig. 4j), confirming the feeding and metabolic effects of high-protein feeding under these conditions. In contrast, the anorectic response to P45 was blunted DMH^Leu-TT^ mice (Fig. 4i) and remarkably the P45 diet did not produce a weight response in these mice (Fig. 4j). Thus, DMH neurons activated by NTS leucine are required for the acute anorectic response to high protein diets, while other pathways likely mediate the sustained anorectic effect of dietary proteins. Unexpectedly, these results indicate that the NTS→DMH leucine-sensing circuit contributes to the metabolic effect of high-protein diets. These results provide a central mechanism for the behavioral and metabolic effects of dietary protein.

## Discussion

Our findings reveal a mechanism through which nutrient sensing in the NTS regulates food-seeking behavior, satiety and long-term energy balance via polysynaptic inhibition of AgRP neurons. We demonstrate that PrRP^NTS^ neurons engage this circuit in response to the detection of the branched-chain amino acid leucine, a signal of dietary protein availability. Silencing of DMH neurons responsive to NTS leucine sensing blunts leucine’s appetite-suppressive effects and dampens the anorexic and weight loss responses to a high protein diet, hence extending the role of this circuits in the behavioral and metabolic responses to dietary proteins.

Our data expand the characterization of the functional diversity of NTS TH neurons to include a subset of neurons expressing PrRP and CTR, projecting to the DMH and modulating feeding initiation and satiety via downstream projections to AgRP neurons. While other neuronal populations (including NTS and PVH neurons) activated by NTS leucine are likely involved in other behavioral, metabolic and neuroendocrine outputs of NTS leucine sensing, the circuit described here is sufficient to entirely explain the appetitive consequences of NTS leucine detection. These results provide support for a model in which specialized neuronal populations regulate specific behavioral outputs. Intriguingly, although PrRP and CTR neurons of the NTS have been shown to project to the PBN (Blouet et al., 2009; Dodd & Luckman, 2014), this site is not activated by NTS leucine, suggesting further functional diversity among these neurons.

Recent work indicates that AgRP neurons integrate various sensory inputs including environmental food-related cues and visceral mechanosensory and nutritional inputs (Bai et al., 2019; Betley et al., 2015; Beutler et al., 2017; Mandelblat-Cerf et al., 2015). Our work extends the integrative capability of AgRP neurons to include brainstem nutrient sensing inputs. Given that visceral vagal afferents terminate in the caudomedial NTS where PrRP/CTR neurons are concentrated, it is likely that intestinal sensory inputs engage the same circuit as NTS leucine to inhibit AgRP neurons. Thus, PrRP^NTS^→ LepR ^DMH^ circuit may be specialized in integrating nutritional cues arising from multiple central and peripheral interoceptors. These sites may be only partially functionally redundant, given the indication that caloric density is the primary information carried by vagal afferents activated by nutrients in the gut (Williams et al., 2016), whereas NTS nutrient sensing surveys nutritional status and post-absorptive nutrient availability. With the ability to monitor the availability of specific nutrients and relay this information to forebrain centers mediating the long-term control of energy balance, PrRP ^NTS^ neurons and their projections to LepR ^DMH^ neurons are well positioned to contribute to the production of nutrient-specific satiety, a well-established behavioral lacking mechanistic characterization (Johnson & Vickers, 1992).

DMH ^LepR^ neurons also process environmental food-related cues in the regulation of AgRP activity, and these may be integrated with former signals as well (Garfield et al., 2016). To our knowledge, these are the only inhibitory inputs to AgRP neurons identified so far. However polysynaptic retrograde tracing from AgRP neurons here revealed additional medullary inputs to AgRP neurons which may extend the inhibitory control of this population. Collectively, this work resolves the mechanisms through which NTS nutrient sensing modulate food-seeking behavior and provides insights into the functional organization feeding-regulatory circuits, creating new opportunity for the treatment of hyperphagic obesity and related metabolic disorders.

## Material and Methods

### Experimental models

All experiments were performed on male mice in accordance with the Animals (Scientific Procedures) Act 1986 and approved by the local animal ethic committees. Mice were obtained from Charles River UK (8-wk old C57/bl6J) or the Jackson Laboratories (*Agrp-ires-cre, Th-cre, NPY-hrGFP*), housed in individually ventilated cages with standard bedding and enrichment, maintained in a humidity-controlled room at 22-24°C on a 12 hr light/dark cycle with ad libitum access to water and standard laboratory chow diet unless otherwise stated. Isocaloric modified diets with varying protein amounts were custom made by Research Diets as per the formulations in Suppl. Table 1. For all experiments using cre reporter lines, we carried the work in hemizygous males or wild-type littermates randomly assigned to experimental groups. For studies performed on wild type mice, weight-matched groups were compared. Before each dietary change, mice were briefly exposed to the new diets to avoid neophobia or other novelty-related response in subsequent experiments.

### Methods details

#### Stereotaxic surgical procedures

Surgical procedures were performed on mice aged 9 to 11-wk old, under isofluorane anesthesia. All animals received Metacam prior to the surgery, 24 hr after surgery and were allowed a 1-week recovery period during which they were acclimatized to injection procedures. Mice were stereotactically implanted with bilateral steel guide cannulae (Plastics One) positioned 1 mm above the ARH (A/P: -1.1 mm, D/V: -4.9 mm, lateral: +/- 0.4 mm from Bregma) or the DMH (A/P: -1.5 mm, D/V: -4 mm, lateral: +/- 0.4 mm from Bregma), or 2mm above the caudomedial NTS (cannula holding bar in a 10° rostro-caudal angle, coordinates relative to occipital suture: A/P +0.5 mm, D/V -3 mm, lateral: +/- 0.4 to midline). Beveled stainless steel injectors (33 gauge) extending 1 mm (for ARH and DMH) and 2mm (for NTS) from the tip of the guide were used for injections. For chronic cannulae implantation, cannula guide was secured in place with Loctite glue and dental cement (Fujicem2). Correct targeting was confirmed histologically postmortem. Mice were allowed 1-wk recovery during which they were handled daily and acclimatised to relevant experimental settings.

#### Viral vectors and injection procedures

For cre-dependent retrograde polysynaptic tracing, we used PRV-introvert, a newly developed version of PRV-Bartha in which retrograde viral propagation and reporter expression are activated only after exposure to cre recombinase with high specificity (Pomeranz et al., 2017), kindly provided by Prof. Jeff Friedman (Rockefeller University). PRV-introvert was prepared as previously described (Pomeranz et al., 2017). Virus stocks were grown and tittered in PK15 cells (7.89 x 10^8^pfu/ml) (ATCC). Viral specificity was tested in vitro in HEK cells transfected with cre and by stereotaxic injection into the ARH of wild-type (n = 5). A total of 25 mice were used to characterize polysynaptic inputs to AgRP neurons. All received 100nl of PRV-Introvert into the ARH and were sacrificed at 0, 24, 48, 72 or 96h) after the injection. These mice rapidly developed symptoms, were closely monitored, provided with hydrogel, mash and a heating pad throughout the postsurgical period, and were killed before reaching 20% of pre-surgical weight loss. Retrograde cre-dependent monosynaptic tracing was performed using AAV8-hSyn-FLEX-TVA-P2A-eGFP-2A-oG (2.82×10^12^ vg/ml, 500nl per side) and the modified rabies strain SADΔG-mCherry(EnvA) (1.1×10^9^ TU/ml, 100nl bilaterally into the ARH 3 weeks later)(Callaway & Luo, 2015; E. J. Kim et al., 2016) both obtained from the Salk Institute Viral Vector Core. A total of 23 mice were used and perfused at different survival times after the rabies injection and up to 2 weeks later. Activity dependent cell labelling and anterograde tracing were performed using the viral construct AAV8-fos-Cre-ERT2-PEST (AAV-fos-CreER, 8.8×10^12^ vg/ml, 300nl per side bilaterally, donated by Prof Deisseroth via the Stanford Virus Core) was combined with AAV8-EF1a-DIO-hChR2(H134R)-mCherry (AAV-DIO-ChR2:mCherry, 1.9×10^13^ vg/ml, 300nl per side, Addgene) or AAV-DJ-CMV-DIO-eGFP-2A-TeNT (AAV-DIO-TeNT, 5.13×10^12^ vg/ml, 300nl per side, Stanford University Neuroscience Gene Vector and Virus Core).

#### NTS Leucine injection and acute food intake assessments

Studies were conducted in a home-cage environment. For NTS leucine injection, mice were food deprived for the 6 hr during the day before receiving a bilateral parenchymal injection of L-leucine (Sigma, 2.1 mM, 50 nl/side, 50 nl/min) or aCSF (R and D) and were either immediately returned to their home cage for food intake analysis or perfused 80/90 min later for histological assessments. For food intake studies, the injection occurred 1h before dark-onset. Mice were refed after the injection and food intake was monitored over various time points after the refeeding. For the meal initiation experiment, digital cameras were used to record the first 30 min feeding respond after the mice received the brain injection and provided with a food pellet. All studies were performed in a crossover randomized manner on age- and weight-matched groups, and at least 4 days elapsed between each brain injection.

#### Activity-dependent induction of cre expression

Mice that received AAV-fos-CreER into the NTS or the DMH and were chronically equipped with a cannula guide targeting the NTS went through a series of induction sessions as follows. Mice received an injection of leucine into the NTS in their home cage as above and 80 minutes later were dosed with the tamoxifen metabolite 4-hydroxytamoxifen (4-PHT, Sigma, 40mg/kg ip) prepared using a formulation described previously (Ye et al., 2016). Mice remained fasted for 4hrs after the 4-HT injection. Each induction sessions were separated by a minimum of 96 hr.

#### Brain perfusion, immunohistochemistry, microscopy and image analysis

Animals were anaesthetized with Dolethal (Vetoquinol UK Ltd) at 1 ml/kg in saline and transcardiacally perfused with 0.1M heparinized PBS followed by 4% paraformaldehyde. Brains were extracted and post-fixed in 4% paraformaldehyde, 30% sucrose for 48 hr at 4°C. Brains were sectioned using a Leica freezing sliding microtome into 5 subsets of 25 microns sections. Antigen retrieval was used for all experiments prior to antibody incubation. Sections were incubated in 10 mM sodium citrate at 80°C for 20 min then washed three times in PBS. Tissue was blocked for 1 hr with 5% normal donkey serum or 5% normal goat serum (Abcam) at room temperature, and incubated at 4°C with primary antibodies against c-fos (1:2000, Synaptic Systems), dsRed (1:1000, Clontech), GFP (1:1000, Abcam), TH (1:200, Immunostar), PrRP (1:1000, Abcam), Ha (1:1000, Cell Signaling Technology) and GFRAL (1:200, Thermo Fisher Scientific). Sections were then mounted on slides and coverslipped with Prolong Diamond (Thermo Fisher Scientific). Sections were imaged using a Zeiss Axio slide scanner with 20x objective or Leica SP8 confocal microscope with the 40x or 63x objectives. Imaging settings remained the same between experimental and control conditions.

Images of tissue sections were digitized, and areas of interest were outlined based on cellular morphology and using the brain atlas of Paxinos and Franklin (Paxinos & Franklin, 2001). Images were analyzed using the ImageJ manual cell counter or Zeiss ZEN 2.3 software. For the analysis of projection coverage to the ARH, Imaris software (Oxford Instruments plc) was used to 3D reconstruct the ARH images stacks acquired by SP8 microscope and analyze the contact areas.

#### Multiplexed FISH with RNAscope

Mice were perfused as described above. Brains were postfixed in 4% PFA solution overnight then cryoprotected in 30% sucrose solution in PBS for up to 24 h. Tissue was covered with optimal cutting temperature (OCT) media then sliced at 16 μm thickness using a Leica CM1950 cryostat directly onto Superfrost Plus slides (ThermoScientific) in an RNase free environment. Slides were then stored at -80°C. Multiplexed fluorescence in situ RNA hybridization (FISH) was performed using RNAscope technology. After epitope retrieval and dehydration, sections on slides were processed for multiplexed FISH using the RNAScope LS Multiplex Assay (Advanced Cell Diagnostics). Samples were first permeabilized with heat in Bond Epitope Retrieval solution 2 (pH 9.0, Leica - AR9640) at 95°C for 2 min, incubated in protease reagent (Advanced Cell Diagnostics) at 42°C for 10 min, and finally treated with hydrogen peroxide for 10 min to inactivate endogenous peroxidases and the protease reagent. Samples were then incubated in z-probe mixtures for 2 h at 42°C and washed 3 times. DNA amplification trees were built through incubations in AMP1 (preamplifier), AMP2 (background reducer), then AMP3 (amplifier) reagents (Leica) for 15-30 min each at 42°C. Between incubations, slides were washed with LS Rinse buffer (Leica). After, samples were incubated in channel-specific horseradish peroxidase (HRP) reagents for 15 min at 42°C, TSA fluorophores for 30 min and HRP blocking reagent for 15 min at 42°C. The following TSA labels were used to visualize z-probes: Cy3 (1:500), FITC (1:500), and Cy5 (1:500) fluorophores (Perkin Elmer).

Brain sections were imaged using a spinning disk Operetta CLS (Perkin Elmer) in confocal mode using a sCMOS camera and a 40x automated-water dispensing objective. Sections were imaged with z stacks at intervals of 1 μm. ROIs included the PVH, DMH, NTS, AP and DMX. Gain and laser power settings remained the same between experimental and control conditions within each experiment. Harmony software (Perkin Elmer) was used to automatically quantify number of labelled RNA molecules (spots) per cell, and number of labelled cells among other metrics.

### Statistical analysis

All data, presented as means ± SEM, have been analyzed using GraphPad Prism 8. For all statistical tests, an α risk of 5% was used to define statistical significance. Dietary and aCSF/Leucine treatments where allocated randomly in weight-matched groups. When possible, we performed within mice comparisons and treatment were delivered in a cross-over manner in weight-matched groups. All kinetics were analyzed using repeated-measures two-way ANOVAs and adjusted with Bonferroni’s post hoc tests. Multiple comparisons were tested with one-way ANOVAs and adjusted with Tukey’s post hoc tests. Single comparisons were made using two-tail Student’s t tests. We used blinding (to mouse genotype, viral treatment or drug delivered) for in vivo experiments and to perform image analysis. Additional statistical details for each experiment can be found on the figures on in the figure legend.

**Supp. Movie 1**: Video recording of a DMH^leu-TT^ mouse following NTS aCSF administration and food presentation (paradigm shown on Fig.1a).

**Supp. Movie 2**: Video recording of a DMH^leu-TT^ mouse following NTS Leu administration and food presentation (paradigm shown on Fig.1a).

**Supplemental Table 1:** Modified diets composition.

| Ingredient | P20 |  | P45 |  |
| --- | --- | --- | --- | --- |
|  | gm |  | gm |  |
| Casein | 233 |  | 510 |  |
| L-Cystine | 3.49 |  | 7.65 |  |
| Corn Starch | 326.6 |  | 156.5 |  |
| Maltodextrin 10 | 150 |  | 75 |  |
| Sucrose | 107.1 |  | 107.1 |  |
| Cellulose | 50 |  | 50 |  |
| Soybean Oil | 88.9 |  | 88.9 |  |
| tBHQ | 0.014 |  | 0.014 |  |
| Mineral Mix S10022G | 0 |  | 0 |  |
| Mineral Mix S10022C | 3.5 |  | 3.5 |  |
| Calcium Carbonate | 10 |  | 12.35 |  |
| Calcium Phosphate, Dibasic | 3.4 |  | 0 |  |
| Potassium Citrate, 1 H2O | 3 |  | 8 |  |
| Potassium Phosphate, Monobasic | 6.31 |  | 0 |  |
| Sodium Chloride | 2.59 |  | 2.59 |  |
| Vitamin Mix V10037 | 10 |  | 10 |  |
| Choline Bitartrate | 2.5 |  | 2.5 |  |
| FD&C Yellow Dye #5 | 0 |  | 0 |  |
| FD&C Red Dye #40 | 0.05 |  | 0 |  |
| FD&C Blue Dye #1 | 0 |  | 0.05 |  |
| Total | 1000.4 |  | 1034.1 |  |
|  | gm | kcal | gm | kcal |
| Protein | 206 | 824.8 | 451 | 1805.4 |
| Carbohydrate | 594 | 2375 | 349 | 1394 |
| Fat | 89 | 800.2 | 89 | 800.2 |
| Fiber | 50 | 0 | 50 | 0 |
| Total | 939 | 4000 | 939 | 4000 |
|  | gm% | kcal% | gm% | kcal% |
| Protein | 21 | 21 | 44 | 45 |
| Carbohydrate | 59 | 59 | 34 | 35 |
| Fat | 9 | 20 | 9 | 20 |

## Acknowledgments

We thank B. Roth and K. Deisseroth for AAV plasmid constructs, E. Callaway for rabies tracing tools, and J. Friedman for sharing pseudorabies virus introvert. We thank Gregory Strachan at the Wellcome Trust-MRC Institute of Metabolic Science for his assistance with image analysis, and Julia Jones and Heather Zecchini at the Cancer Research UK Cambridge Institute for their assistance with RNAscope staining and imaging. This work was supported by the Medical Research Council (MR/N003276/1), the Medical Research Council Metabolic Disease Unit, and the Wellcome Trust Strategic Award for the MRL Disease Model Core and Imaging facilities (MRC_MC_UU_12012/5 and 100574/Z/12/Z).

## Author Contributions

AHT performed experiments, data analysis and drafted the manuscript. DN performed experiments, data analysis. TD and HS performed experiments. CB designed and performed experiments, data analysis and prepared the manuscript.

## Competing Interests

The authors declare no competing interests.

## Materials and correspondence

Correspondence and material requests should be addressed to Clemence Blouet.

## Notes

### Competing Interest Statement

The authors have declared no competing interest.

